# Furin Inhibition Protects Against Acute Lung Injury in a Mouse Model of Pseudomonas Aeruginosa Infection

**DOI:** 10.1101/2025.09.25.678606

**Authors:** Olivier Bernard, Michael Kwon, Mark R Looney, Mélia Magnen, Michelle A Yu

## Abstract

*Pseudomonas aeruginosa* (PA) is responsible for significant morbidity and mortality particularly in patients with chronic lung diseases such as chronic obstructive pulmonary disease (COPD), bronchiectasis, cystic fibrosis (CF) as well as hospital-acquired pneumonia (HAP) and ventilator-associated pneumonia (VAP). The rise of antibiotic-resistant PA strains necessitates alternative treatment strategies. Among the different toxins secreted by PA, Exotoxin-A (Exo-A) becomes cytotoxic when cleaved by furin. This study investigates the therapeutic potential of furin inhibitor BOS-318 in mitigating acute lung injury induced by Exo-A and PA infection. Furin inhibition significantly improved survival rates and reduced lung injury in mouse pneumonia models using Exo-A and PA103. Additionally, BOS-318 accelerated bacterial clearance in vivo, and increased phagocytosis by alveolar macrophages. Bulk RNA-seq immune profiling revealed modulation of the natural killer (NK) cell signaling pathway possibly due to a decrease in NK recruitment, suggesting a role of furin in shaping the immune response. Overall, our findings demonstrate that furin inhibition protects against PA-induced acute lung injury and hastens bacterial clearance. These results are the first to characterize furin inhibition in animal models and supports its potential use as an adjunctive therapeutic strategy for treating PA infections.

## INTRODUCTION

Pseudomonas aeruginosa (PA), an opportunistic gram-negative bacterial pathogen, is a common cause of morbidity and mortality in hospital-acquired pneumonia (HAP) and ventilator-associated pneumonia (VAP). immunocompromised patients, and those with structural lung disease such as COPD and CF (1, 2). Despite recent therapeutic advances, PA infection remains a significant challenge in CF. The prevalence of PA in adults with CF was estimated in 2021 at 34.2% and is associated with significant morbidity such that upon first colonization, extensive antibiotic courses are used in an attempt at eradication (3, 4). PA leads to increased lung function decline, pulmonary exacerbations, and mortality from terminal respiratory failure due to inflammation and airway damage. PA’s ability to form biofilms complicates treatment by hindering antibiotic penetration, reducing drug efficacy, and promoting antibiotic resistance. Despite effective CFTR modulator therapies, clonal lineages persist in CF patients, particularly in those with advanced lung disease. Therefore, new therapies that assist eradication of PA and decrease PA-associated lung inflammation are needed in CF, and these advances could also greatly benefit COPD, HAP, and VAP patient populations (5). PA employs both intrinsic and acquired resistance mechanisms, making successful eradication increasingly challenging (6). Through the secretion of virulence factors, PA causes cellular damage, impairs the immune response, and enhances its own survival (6). Among these factors, exotoxin-A (Exo-A) is one of the most prevalent virulence factors in resistant clinical isolates of PA and mediates virulence by inhibiting protein synthesis in host cells as well as delivering effector proteins directly into host cells, disrupting cellular processes and contributing to infection (7). As a result, higher titers of IgG antibody to Exo-A are associated with mortality in patients with CF (8). After being secreted by PA, Exo-A is internalized and cleaved by a proprotein convertase, PCSK3 also known as furin. Its toxic effect arises from its ability to inactivate cytosolic elongation factor 2, thereby inhibiting protein biosynthesis leading to cell death (9-11). Exo-A has been shown to impair natural killer (NK) cell cytotoxicity functions (12). Moreover, another virulence factor secreted by PA, Exo-T has been shown to increase numbers of NK cells in the lung tissue during PA-induced pneumonia (13). BOS-318 is an exceptionally potent and highly selective furin inhibitor, which prevents Exo-A-induced cell toxicity in human primary epithelial cells (14).

For the first time, we report that BOS-318 modulates Exo-A and PA103-induced lung injury in animal models. Remarkably, inhibiting furin in our mouse models increased survival and reduced lung injury. In the live bacteria model, bacterial clearance was accelerated, facilitated by enhanced alveolar macrophage phagocytosis. Interestingly, furin inhibition also modulated NK signaling, reducing inflammation. Together, our findings support furin inhibition as a therapeutic strategy to treat *Pseudomonas aeruginosa* infection.

## METHODS

### Care and use of mice

Mice were housed under pathogen-free conditions at the UCSF Laboratory Animal Research Center and all experiments conformed to ethical principles and guidelines approved by UCSF Institutional Animal Care and Use Committee. Mice were provided with water and food *ad libitum*, and supplemental water gel packs and food were placed in the cage during experiments. 10 to 12-week-old C57BL/6J wild-type male mice were obtained from the Jackson Laboratory for lung infection studies. Our study exclusively examined male mice. It is unknown whether the findings are relevant for female mice.

### Bacterial cultures

PA103 was kept at −80 deg-C and streaked onto plates overnight after which two single colonies were selected to inoculate the subculture and promote purity using a previously published protocol (15). For each experiment, 2 single colonies of *Pseudomonas aeruginosa* strain PA103 were grown overnight in tryptic soy broth (TSB) at 37ºC, shaking at 200 rpm. The next day, a sub-culture of bacteria broth was done at a dilution of 1:20 and harvested at the mid-log phase [optical density (OD) 0.50 at 600 nm], (Genesys 10S UV-Vis, Thermo scientific). Bacterial culture was spun 15min at 2,700 g, 4C and resuspended in ice-cold PBS.

### BOS-318

For intraperitoneal (i.p.) administration, BOS-318 (HY-147140, MCE) was resuspended in 10% DMSO, 90% corn oil at [10mM]. For oropharyngeal administration (o.a.), BOS-318 was resuspended in DMSO at [8.74mM].

### Lung injury models

A solution of 100 µL PBS + 100 µL of 10% DMSO / 90% corn oil (= vehicle) or BOS-318 was injected i.p. 1h prior to o.a. Mice were then anesthetized with isoflurane 5%, then 50 µL of a mixture composed of 40 µL PBS + 5 µL DMSO or BOS-318 + 5 µL of Exotoxin-A [= 0.25µg] (Sigma) or PA103 was delivered via o.a. Mice were euthanized at different time points to collect either lungs or broncho-alveolar lavage (BAL). Mice lungs were lavaged with 1 mL of PBS. Bacterial colony forming units (CFU) of lung homogenates were quantified by serial dilutions on Pseudomonas Isolation Agar (PIA) plates in duplicate and averaged. Colonies present on plates were manually counted after overnight culture at 37C, in dilutions where 10-100 colonies could be counted. BAL was centrifuged 5min at 300g, 4C. The supernatant was collected and stored at −80C for further analysis. BAL cells were incubated for 30 min with APC-Cy7-zombie NIR (50-604-714, Biolegend) at RT in the dark, centrifuged, then fixed for 20 min with Cytofix (BDB554722, BD), then incubated with anti-mouse CD16/32 (clone 2.4G2 BE307, BioXCell) for 10 min to block non-specific binding. Samples were then incubated for 30 min with an appropriate antibody cocktail. Antibody reagents include: BV711-CD45 (clone 30F11, BDB563709, BD), PE-CD11b (clone M1/70, 101208, Biolegend), PE-Cy7-Ly6G, AF488-Ly6C (clone HK1.4, 50-112-4604 Invitrogen), Siglec-F BV510 (clone E50-2440 BDB740158, BD). Counting beads were used for BAL analysis to count absolute numbers of cells in the sample (C36950, Thermo Fisher). Beads were used for compensation (50-112-9040, BD). Cells were recorded using a LSR II flow analyzer (BD) and analyzed with FlowJo (BD). Alveolar macrophages are CD45+ Siglec F+ F4/80+. Neutrophils are CD45+ Siglec F-F4/80-Ly6G+ Ly6C+. Monocytes are CD45+ Siglec F-F4/80-Ly6G-Ly6C+.

### Flow cytometry-based phagocytosis assay

BAL from untreated WT mice was centrifuged for 5 min x 300g, then the cell pellet was resuspended in 200 µ**L** of DMEM 10% (11-965-092, Gibco) FCS 1% Penicillin/streptomycin (15140122, Gibco) and split in two wells in a 96-well plate (Falcon) with DMSO or BOS-318 at [1 µM] final concentration for 2h. Media was then replaced and *E. Coli* bioparticles conjugated with the pH-sensitive fluorescent dye pHrodo (P35366, Invitrogen) were added at [0.1 mg/ml]. Following an incubation of 1h30, cells were detached with TrypLE (12604013, Gibco), centrifuged and resuspend in MACS before flow analysis. On selected experiments, purity was assessed using antibody staining and flow cytometry.

### BAL analysis

The total protein content in the BAL was determined by BCA Protein Assay Kit (23225, Thermo scientific). BAL albumin concentration was quantified using Mouse Albumin ELISA Kit (E99-134, Fortis Life Sciences). Cytokines were measured using R&D ELISA kits (MCP-1 DY47905, MIP-2 DY45205, KC DY453). Glucose-6-phosphate dehydrogenase (G6PD) level was measured using CyQUANT™ Cytotoxicity Assay kit (V23111, Invitrogen). Sodium levels were determined in BAL according to manufacturer instructions (abx298875, Abbexa)(16).

### Histology

Lungs were fixed in 4% (vol/vol) PFA, embedded in paraffin, sectioned at 4 µm, and stained with H&E, TUNEL (G7130, Promega) or NK1.1 (clone E6Y9G, cell signaling) by immunohistochemistry (Histowiz). TUNEL positive areas, i.e. positive stained areas vs total tissue areas, of the whole lower part of the right lobe (analysed lobe: 1115651 px^2^ +/-250489 px^2^) were counted using ImageJ software (17).

### Bulk-RNAseq

Total RNA was extracted from whole mouse lung using RNAeasy kit (74104, Qiagen), quantified by NanoDrop (Thermo) then sent for sequencing by UCSF Genomic CoLabs. Samples were validated for QC then sequenced using NovaSeqX sequencer. 50 bp paired-end reads were acquired on a NovaSeq X at the UCSF Center for Advanced Technology. FastQC was used to examine the quality of raw sequencing reads. Sequences were aligned to the GRCm38 genome and gene counts obtained with STAR. Raw gene counts were used for differential gene expression analysis with DEseq2. Cutoffs for DEGs were set at a log2 fold change >1 and an adjusted P < 0.05. DEGs between BOS-318 and vehicle groups were subjected to GO analysis (https://geneontology.org/). The heatmap was generated with pheatmap. KEGG 2019 mouse table was generated using genes with FDR < 0.01 in gene ontology, and the first 10 pathways with the lowest adjusted p-value were displayed with ggplot in R. Volcano plots were generated with EnhancedVolcano in R (RStudio 2025.05.0+49).

### Statistical Analyses

Comparisons between two groups were done with relevant paired or unpaired *t*-test or Mann–Whitney *U* test when data were not normally distributed. Comparisons of more than two groups were made with ANOVA (followed by Tukey’s or Dunnett’s multiple comparisons tests) or Kruskal–Wallis (followed by Dunn’s multiple comparisons test). In cases of missing data, a mixed-effects model was used. Survival analysis was done with log-rank test with test for trend where appropriate. *P* < 0.05 was considered statistically significant. Statistical analysis and graph production were done with GraphPad Prism 10 (GraphPad, La Jolla, CA).

### Study Approval

All animal experiments were approved by the Institutional Animal Care and Use Committee at UCSF.

### Data availability

Bulk RNA_seq data and raw data have been provided as a supporting data file.

## RESULTS

### Administration of a furin inhibitor protects mice from Exotoxin-A lung injury

Previously, Douglas *et al*. demonstrated that a new furin inhibitor, BOS-318, prevents exotoxin-A induced cytotoxicity in explanted human bronchial epithelial cells (14), yet whether BOS-318 affects baseline inflammation *in vivo* is unknown. Here, we investigated whether furin inhibition prevents exotoxin-A induced lung injury in exogenous toxin and live bacterial mouse models of lung injury. To characterize safety, BOS-318 or vehicle (DMSO) was administered by o.a. at day 0. Another treatment was done by i.p. one hour prior to o.a. and every 24h until 72h, and mice were monitored and sacrificed at 96h (Supplemental Figure 1A). No differences were observed in total immune cell population, neutrophils and monocytes counts in the BAL between groups (Supplemental Figure 1B-D). Furin inhibition has been shown to prevent cleavage of newly expressed α and γ subunits of ENaC *in vitro* (18) potentially affecting its activity, and sodium concentration in the respiratory tract. Here, the sodium concentration in the BAL remained unchanged after BOS-318 treatment (Supplemental Figure 1E). Therefore, BOS-318 does not affect sodium homeostasis or induce lung inflammation.

We then tested the inhibitor using an Exo-A-induced lung injury model. As described above, mice were pretreated with BOS-318 or vehicle i.p. then administered Exo-A + BOS-318 or vehicle by o.a. (Fig 1A). Body temperature was significantly decreased in the vehicle group at 48h after Exo-A, while mice treated with the furin inhibitor remained relatively stable. Both groups had lower body temperatures, but the BOS-318 treated mice were significantly less hypothermic (Fig 1B). Mice treated with the vehicle also lost more weight than the BOS-318 treated group after 48h and 72h (Fig 1C). Moreover, mortality at 96h in the vehicle group was significantly greater with only 12.5% survivorship in the vehicle group compared to 78% in the treatment arm (Fig 1D). Taken together, BOS-318 conveyed a survival advantage in the Exo-A mouse model. To test for damage to the alveolar-capillary barrier, we measured total protein in the BAL, which was higher in the control group, suggesting a protective role of BOS-318 in Exo-A induced lung damage and permeability (Fig 1E). Overall, these results demonstrate that BOS-318, reduces Exo-A induced lung injury and mortality in mice.

**Fig 1:**
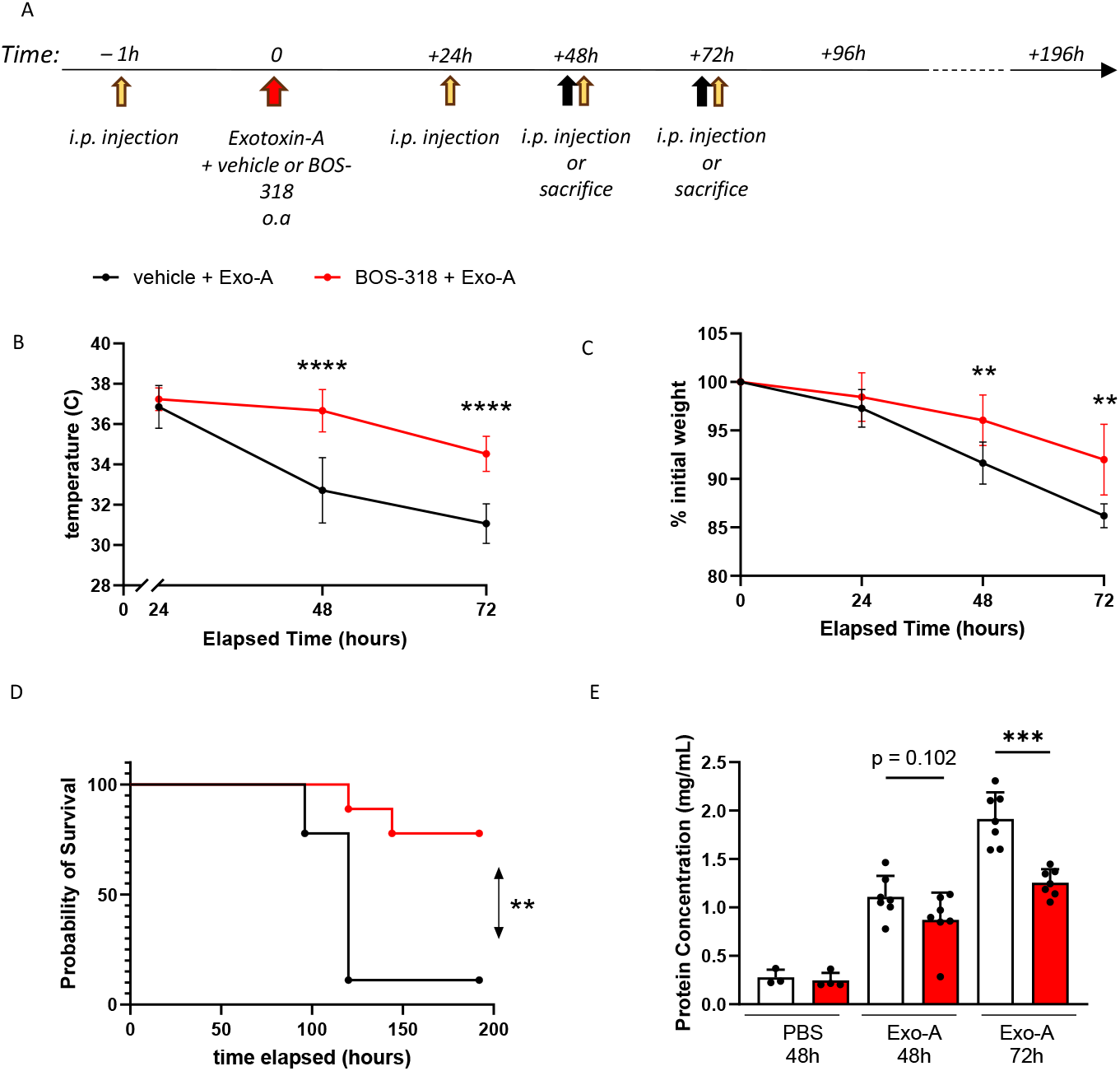
Furin inhibition prevents Exotoxin-A-induced acute lung injury in vivo. (A) schematic depicting experimental procedure. Mice were injected intra-peritoneally (i.p.) with 200ul furin inhibitor BOS-318 [5mM] final or vehicle one hour prior administration oropharyngeal aspiration (o.a.) with 0.25ug of Exotoxin-A with 5ul of BOS-318 [0.874mM] final or DMSO as a control. (B, C) Temperature and % weight loss. **: p < 0.01, ****: p < 0.0001 with a 2-way ANOVA with post-hoc Tukey’s multiple comparisons test. (D) Survival curve. n = 9 mice per group, **: p = 0.0025 with a log-rank test. (E) Bronchoalveolar lavage (BAL) protein concentration.*** p, 0.0001 with unpaired t-test (n = 3-7 per group).

### Administration of a furin inhibitor decreases *Pseudomonas aeruginosa*-induced lung injury and mortality

Exo-A is one of several extracellular toxins secreted by PA, which can induce leukopenia, acidosis, circulatory collapse, liver necrosis, pulmonary edema, hemorrhage, and acute tubular necrosis of the kidneys. Thus, we pursued a live bacterial model to evaluate the effects of BOS-318 against the full spectrum of PA-induced virulence. The [Centers for Disease Control and Prevention](https://www.cdc.gov/index.html) has declared that Pseudomonas strain PA103 is a major concern in healthcare settings due to its ability to cause outbreaks and its high level of antibiotic resistance. Moreover, PA103, produces significantly higher levels of Exo-A than the prototypic strain PAO1 (19). Using our protocol described above, mice were pretreated i.p. prior to infection with BOS-318 or vehicle (Fig 2A). In the survival experiment, mice were challenged with o.a. of 2×10^5^ CFU PA103. In the BOS-318 treated group, body temperature was significantly higher as early as 4h after PA103 injection (Fig 2B). Body weight was lower in the control group at 10h compared to the BOS-318 treated group (Fig 2C). All mice from the control group died within 48h, while to the BOS-318 group exhibited 40% survival (Fig 2D).

**Fig 2:**
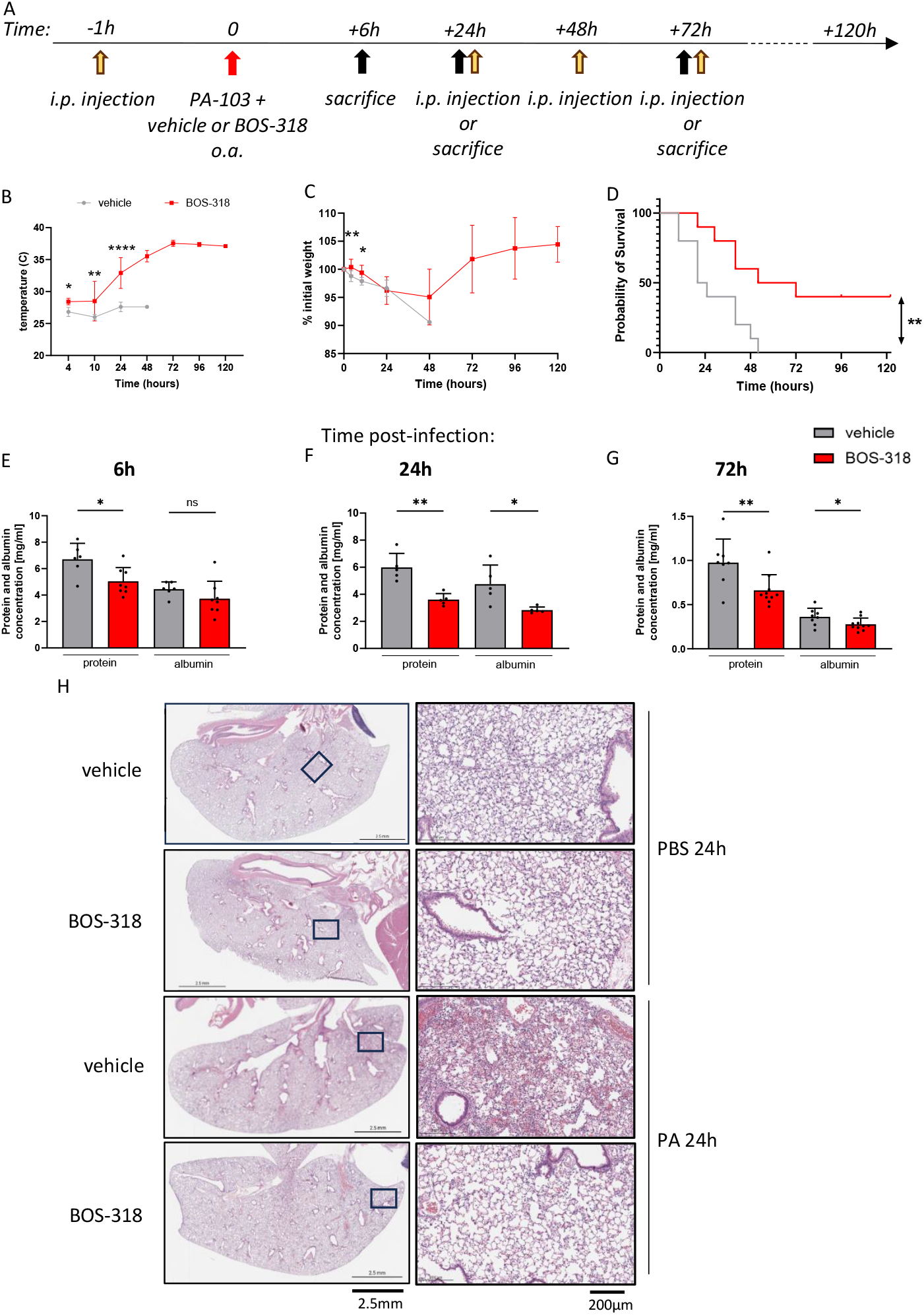
Furin inhibition decreases Pseudomonas Aeruginosa-induced lung injury and mortality. (A) schematic depicting experimental procedure. Mice were injected intra-peritoneally (i.p.) with 200ul furin inhibitor BOS-318 [5mM] final or vehicle one hour prior oropharyngeal aspiration (o.a.) with 2×10^5 CFU of PA103 for survival experiment or 1×10^5 CFU for animals aiming to be sacrificed in 45ul with 5ul of BOS-318 [0.874mM] final or DMSO as a control. (B, C)Temperature and percentage of weight loss. * P< 0.05, ** P< 0.01, *** P< 0.001, **** P< 0.0001 Mixed-effect analysis with a Tukey’s multiple comparisons test. (D) Survival curve (n = 10 mice per group), **: p < 0.01 with a log-rank test. (E-G) Bronchoalveolar lavage protein and albumin concentration at 6h, 24h and 72h post-infection. * p, 0.05 with unpaired t-test (n = 7-8 per group). (H) representative microphotograph of mouse left lung 24h post-infection by H&E. Scale 2.5mm (left) and 200um (right). N = 2-6/group.

Next, we studied furin inhibition in a non-lethal infection model to characterize long-term effects of immune responses and tissue damage due to PA. Mice were infected with 1×10^5^ CFU. BAL protein levels were elevated as soon as 6h post-infection and remained elevated at 72h (Fig 2E-G). The BOS-318-treated group had lower protein levels in BAL compared to the vehicle group at all times, indicative of reduced lung permeability. Similar differences were seen in BAL albumin content at 24h, suggesting that furin inhibition protected against vascular leak. At 24h, PA103 infection led to a dense inflammatory infiltrate characterized by alveolar wall edema and neutrophilic infiltration into the lung parenchyma, which was significantly reduced in the BOS-318 group (Fig 2H).

### Furin inhibitor reduces *Pseudomonas aeruginosa*-associated cytotoxicity

Furin is known to activate Exo-A, enhancing PA-induced cell death in the lung. Furin inhibition using BOS-318 has been shown to be effective in vitro to prevent PA-related toxin Exo-A induced cell toxicity (14). To investigate whether furin inhibition can rescue the lung epithelium from PA103 associated toxicity *in vivo*, we used TUNEL staining detecting DNA fragmentation to label cell death. Furin inhibition led to decreased TUNEL staining at 24h (p =0.0135) (Fig 3A, B). As a second method to assess cell death in lung airspaces, we measured glucose-6-phosphate dehydrogenase (G6PD) activity in BAL. Damaged and dying cells release G6PD, a cytosolic enzyme leaking from cells when plasma membrane integrity is compromised (20). G6PD activity in BAL was significantly decreased in the BOS-318 group at 24h then 72h (Fig 3C-E), which correlates with albumin infiltration in the airspaces (Fig 2E-G). Overall, those results indicate that furin inhibition ameliorates PA-induced cell death and protects the lung epithelium.

**Fig 3:**
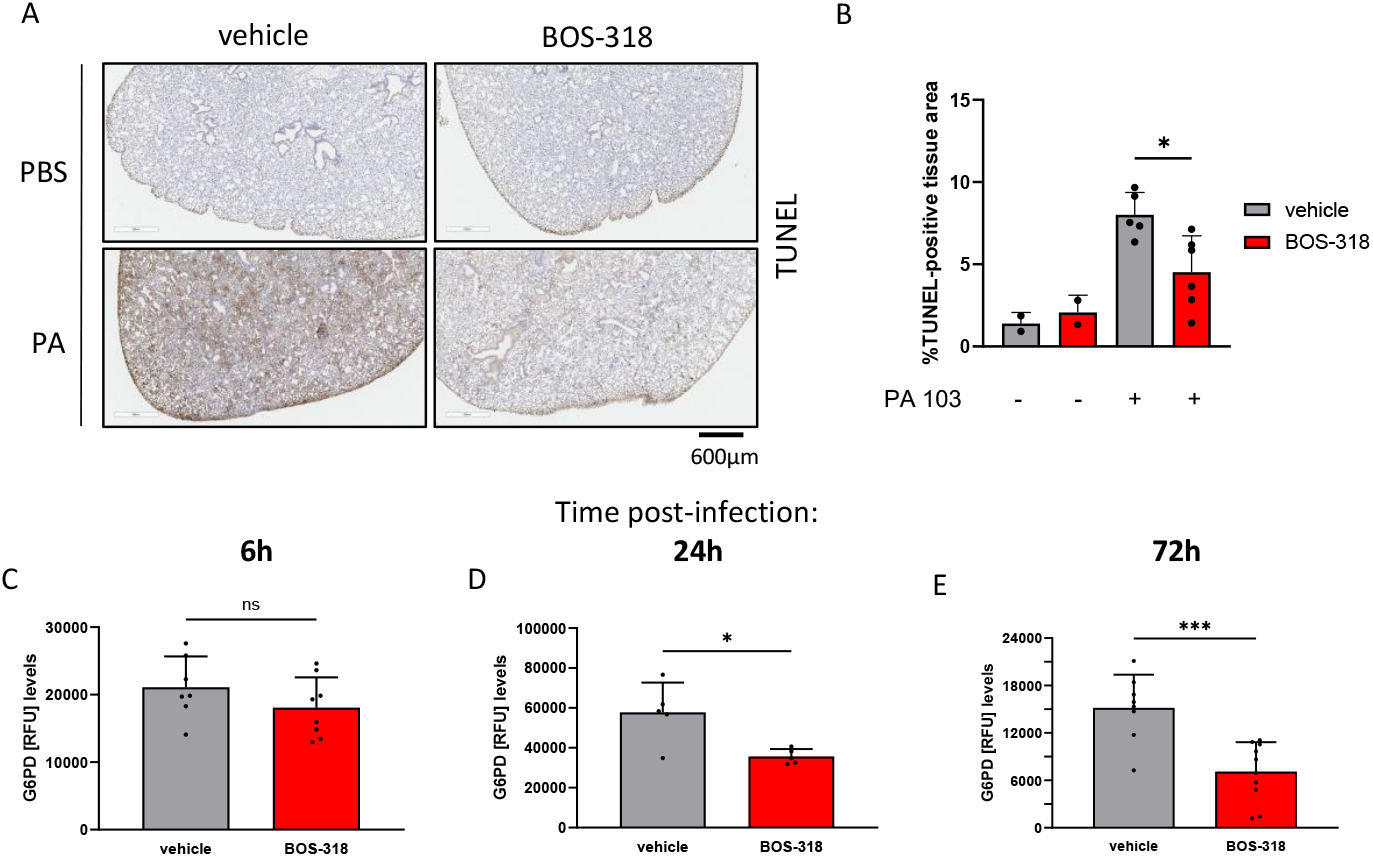
Furin inhibitor reduces Pseudomonas Aeruginosa-associated cytotoxicity. (A) Lungs from mice were collected 24h post-infection and stained for TUNEL. Scale 600um. (B) Percentage of TUNEL-positive areas calculated in the left lobe using high-resolution photomicrographs. n = 2-6 mice/group. * p < 0.05 using unpaired t-test. (C-E) G6PD enzyme activity assayed in Bronchoalveolar lavage (BAL) realized at 6h, 24h and 72h post-infection. n = 5-10 mice/group. * p < 0.05, *** p < 0.0001 using unpaired t-test.

### Furin inhibition improves bacterial clearance and neutrophilic inflammation

The lung epithelium provides the first line of defense against respiratory pathogens by releasing inflammatory mediators that precipitate immune responses. We demonstrated that furin inhibition improves PA-associated cell death, which we postulated occurs by modulating inflammatory responses. We characterized major chemokines in the BAL in our mouse models. The murine chemokines KC (CXCL1) and MIP-2 (CXCL2) are the major chemoattractants responsible for neutrophil recruitment (Fig 4A-C). MIP-2 was significantly (and KC trending to be) less secreted in the BOS-318 group as early as 6h. Another chemokine, CCL2 also called monocyte chemoattractant protein-1 (MCP-1) was also decreased at 6h (Fig 4A-C), which suggests that BOS-318 reduces proinflammatory mediator secretion following PA103 infection. To explore the effects on immune cell recruitment to the lung, BAL samples were analyzed by flow cytometry. CD45+ cells in BAL did not differ at 6h between groups, however BOS-318 treatment led to less total WBC counts, which were predominantly neutrophilic at 24h and 72h (Fig 4D-F). In bacterial pneumonia, neutrophils drive a pro-inflammatory process called NETosis, or neutrophil extracellular trap (NET) formation. This web-like structure induces tissue damage and is targeted by therapies such as recombinant human DNase, or Pulmozyme, in human CF patients (21, 22). BAL NETs were significantly lower in the BOS-318 group at 24h and 72h compared to control (Fig 4J-L), mirroring the decreased neutrophil counts. Resident immune cells in the lung, such as alveolar macrophages (AMs), are responsible for pathogen detection and phagocytosis. AM counts was higher in the BOS-318 group in early time points (6h and 24h) and were more abundant at 24h and 72h (Fig 4G-I).

**Fig 4:**
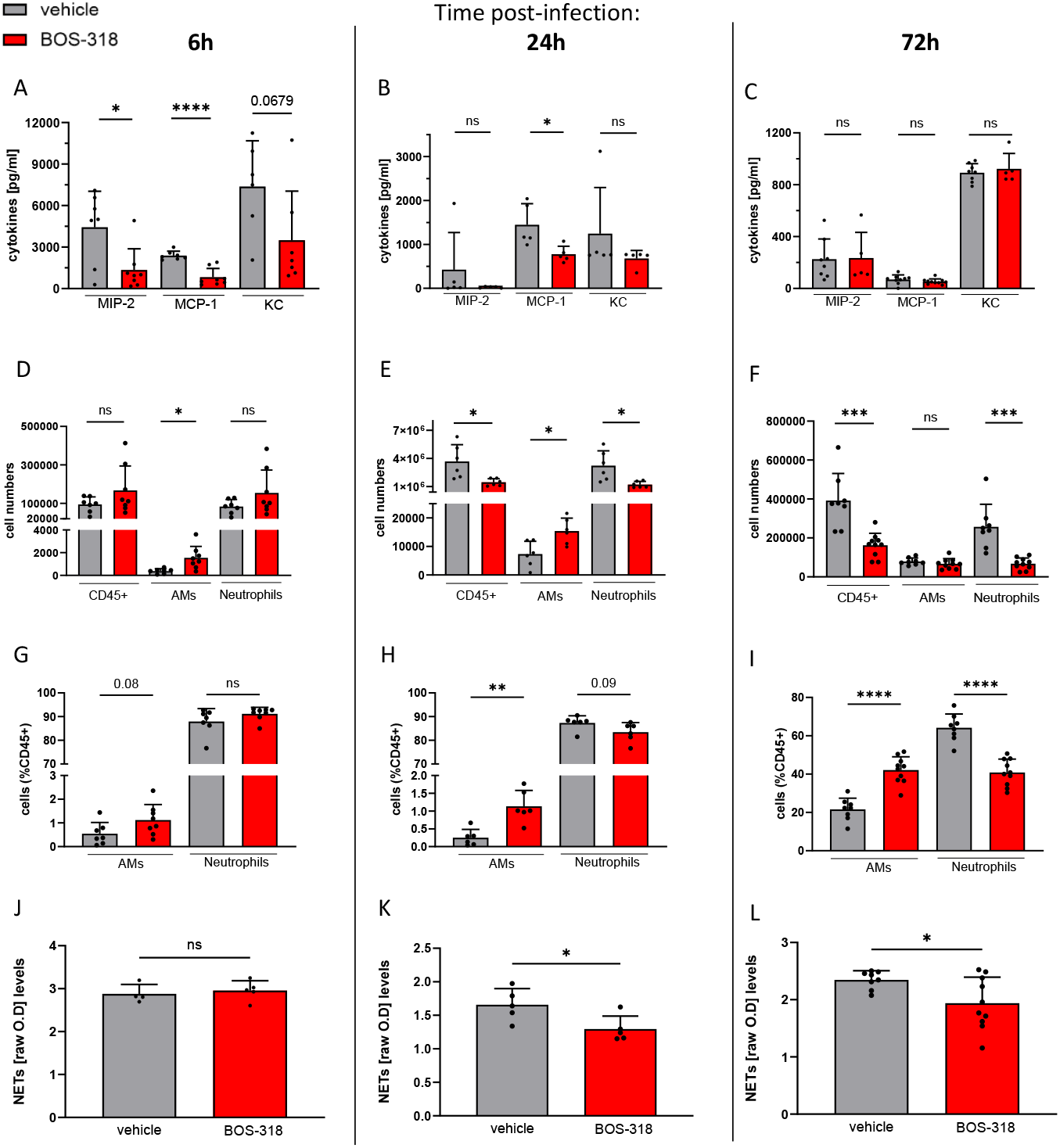
Furin inhibition improves bacterial clearance and neutrophilic inflammation. (A-C) BAL cytokine (MIP-2, MCP-1, KC) expression and neutrophil-elastase-DNA levels at 6h, 24h and 72h post-infection. (D-I) BAL analysis of CD45+, alveolar macrophages and neutrophils numbers or percentage determined by flow cytometry using counting beads. n = 5-8 mice/group. (J-L) Neutrophil extracellular traps secretion in BAL. n = 4-10 mice/group. * p < 0.05, ** p < 0.01, ** p < 0.001, **** p < 0.0001

As immune cells are paramount for bacterial clearance, we explored the effect of BOS-318 treatment on bacterial load. CFU counts in whole lungs at 2h showed that the bacterial burden was unchanged between our two groups, indicating that the inhibitor does not directly affect very early bacteria growth or survival (Fig 5A). As early as 6h p.i., PA burden was significantly decreased in the BOS-318 group, this effect persisted at 24h. Interestingly, the decrease of bacterial burden at 6 h post-infection does not coincide with increased neutrophil counts in the BAL. However, there are more alveolar macrophages numbers in the BOS-318 group. Alveolar macrophages have been described to contribute to PA clearance (23), hence, we explored the effect of furin-inhibition on alveolar macrophages phagocytosis. To do so, mouse alveolar macrophages were pre-treated with BOS-318 before exposure to *E. coli* bioparticles conjugated with pHrodo. The fluorescence of the pHrodo dye dramatically increases as pH decreases from neutral to acidic in phagolysosomes. Hence, phagocytosis is detected by fluorescence detection using flow cytometry (Fig 5B). BOS-318 pre-treated alveolar macrophages displayed increased phagocytosis capabilities (Fig 5C). This effect was specific to alveolar macrophages as furin inhibition did not modulate neutrophil phagocytosis activity (data not shown). These results, suggest that furin inhibition accelerated bacterial clearance by increasing alveolar macrophage phagocytosis activity.

**Fig 5:**
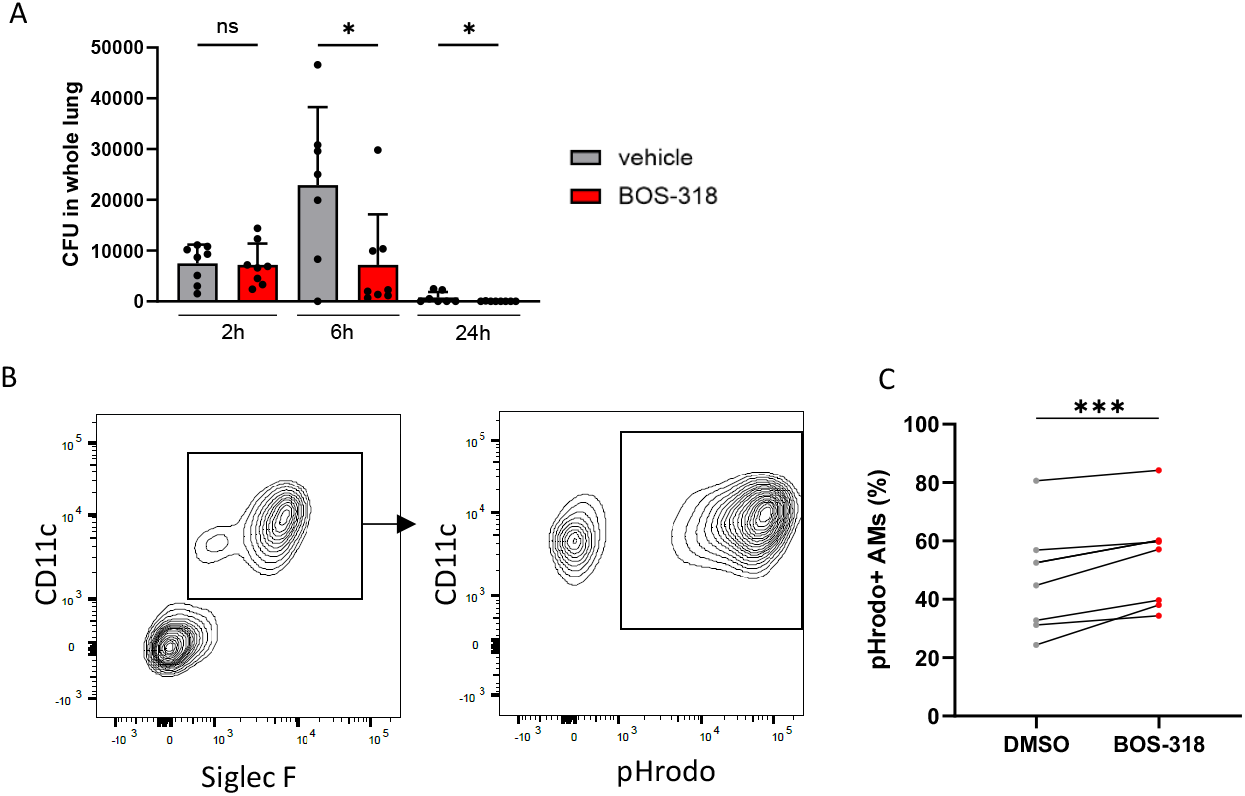
Furin inhibition fastens bacterial clearance and reduce neutrophil infiltration. (A) Mouse lungs were collected at 2h, 6h or 24h post-infection and CFUs are counted (one point per mouse). * p < 0.05 with unpaired t-test (n = 7-8 mice per group). (B,C) Mouse-isolated alveolar macrophages incubated with DMSO or BOS-318 for 2h then media was replaced by pHrodo for 1h30 before recording phagocytosis by flow cytometry. *** p < 0.001 with paired t-test (n = 7 mice).

**Fig 6:**
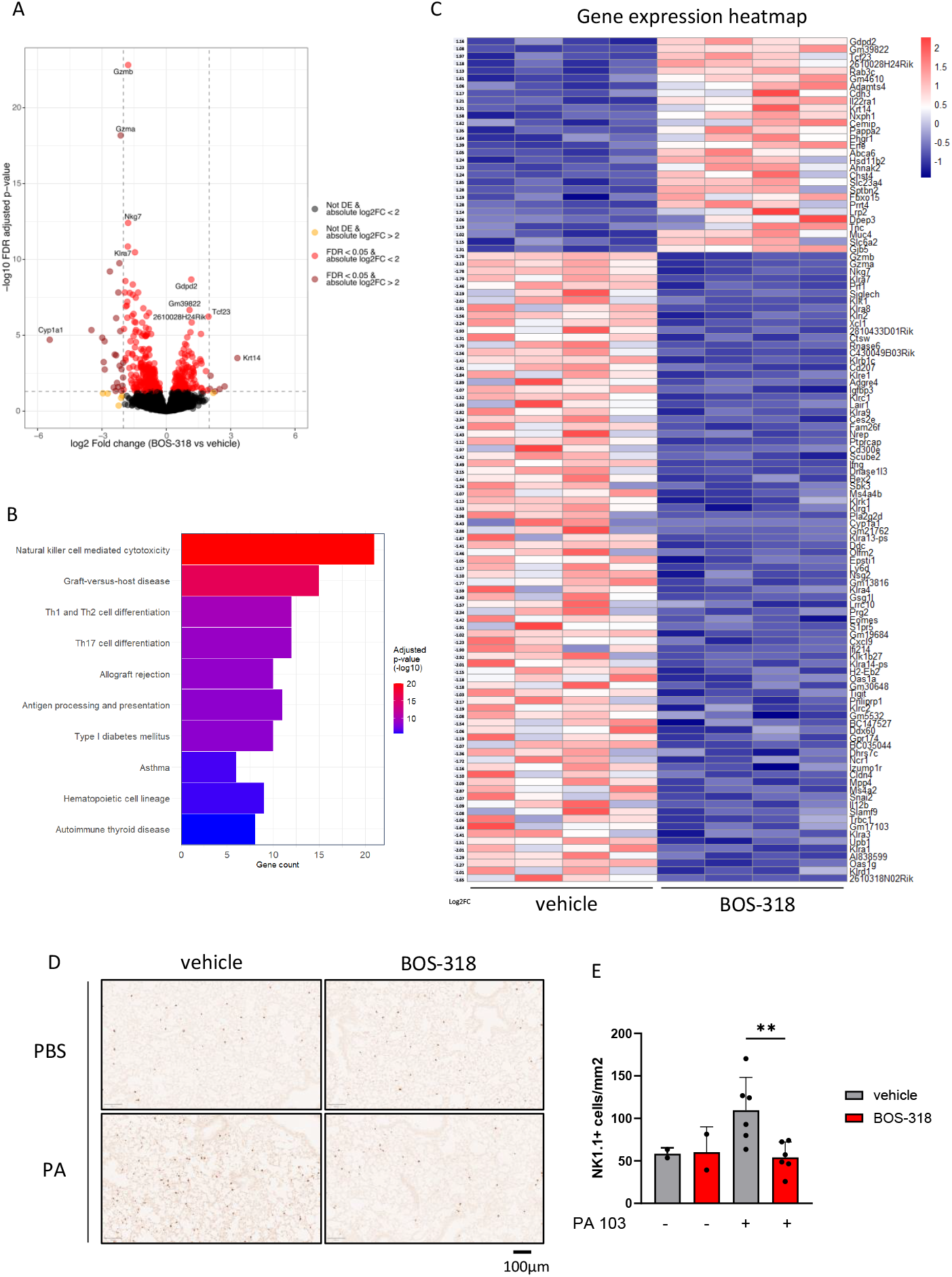
Transcriptomic analysis of PA-infected lungs reveals a downregulation of natural killer cell-related genes. Bulk RNA_seq was performed on whole Lungs 6h post-infection. n = 4 mice/group. (A) Volcano plot displaying genes with FDR < 0.05. (B) KEGG mouse pathways with DFR < 0.01. (C) Heatmap displaying genes with FDR < 0.01. (D-E) Lungs from mice were collected 24h post-infection and stained for NK1.1. Scale 100µm n = 2-6 mice/group. ** p < 0.01

### Transcriptomic analysis of PA-infected lungs reveals a downregulation of natural killer cell-related genes

Furin has pleiotropic effects by processing a wide range of precursor proteins into their active forms, influencing various biological processes. To identify pathways or molecular targets modulated by furin during early PA infection, we performed bulk RNA-seq on mouse lung samples at 6h post-infection (Fig 5A-C). Few genes were upregulated in the BOS-318 group (Fig 5A), but no distinct pathways were highlighted using gene ontology enrichment analysis. Interestingly, the most significantly downregulated pathway in the BOS-318 treated group highlights a NK cell population signature and thus, can be interpreted by a lower accumulation of NK cells in the lung tissue (Fig 5B). NK-related top genes downregulated in the BOS-318 group are shown in the volcano plot and the heatmap (Fig 5A, C), including granzymes A, B, Natural Killer Cell Granule Protein 7 (NKG7), killer cell lectin-like receptor subfamily A member 7 (klra7) or perforin. To decipher whether furin inhibition modulates actual signaling molecules or cell number, we then tested for NK cells in the lung tissue. We found that the influx of NK cells at 24h post infection was reversed by furin-inhibition. Our data show that furin inhibition limits NK cells infiltration in the lung following PA-infection.

## DISCUSSION

In this study, we provide the first characterization of furin inhibition using selective BOS-318 in animal models, advancing an approach to modulate the host-pathogen immune response in bacterial pneumonia. Our major findings include: 1) BOS-318 attenuates Exo-A-induced lung permeability and mortality; 2) BOS-318 improves survival by reducing lung inflammation and tissue damage, evidenced by decreased protein leakage in BAL, decreased neutrophilic infiltration and NETosis, and a preservation of alveolar macrophages; 3) BOS-318 enhances phagocytosis by alveolar macrophages *in vitro* and bacterial clearance *in vivo*. We also showed a preventative effect of BOS-318 on NK cell infiltration. Together, these observations support that furin inhibition is an effective therapeutic approach to mitigate *PA*-mediated pulmonary damage.

The cytotoxic domain of Exo-A has even been harnessed in immunotherapy approaches, where it is fused to tumor-related antigens, in order to penetrate cancer cell membranes and inhibit protein synthesis (24), highlighting its potent biological activity. Indeed, targeting this toxin secreted by *PA* could dampen its virulence and reduce the severity of acute infections and chronic inflammation. Several non-specific furin inhibitors have been developed, including Hexa-D-arginine, which show protection in a Exo-A mouse model, however, the lack of specificity and potency of the inhibitors have resulted in these agents stalling in the preclinical pipeline (25, 26) (27). Our findings are in concordance with previous work by Douglas et al. (14), which demonstrated that BOS-318 potently and selectively inhibited furin-mediated cleavage of Exo-A *in vitro*, thereby preventing its cytotoxic effects. By extending these observations to live bacteria animal models, our study advances the therapeutic effects of furin inhibition in a pathologic relevant context. Interestingly, Amirmozafari et al. observed in human subjects that expression of *ToxA*, an exotoxin-A related gene, correlates with antibiotic resistance (12). One compelling, unanswered question is whether BOS-318 administered synergistically with antibiotics would be effective in treating *PA*. If so, furin inhibition could potentiate current antibiotic susceptibility and thereby reduce antibiotic-resistance in bacteria (28).

CF and COPD share a phenotype of chronic neutrophilic lung inflammation, punctuated by acute bacterial infections, giving them an unusual “acute-on-chronic” inflammatory phenotype, which stands out from other lung diseases (29). Both diseases result in structural damage in which *PA* colonization and infection increase mortality, yet this mechanism is not well understood. Definitive bacterial clearance from the lungs is key to preventing the unrelenting inflammation that leads to lung injury. Our data is the first to show that furin inhibition promotes bacterial clearance in animal models, which we propose is due to decreased neutrophilic infiltration and preservation of alveolar macrophages, resulting in enhanced phagocytic clearance of bacteria. The density of alveolar macrophages has been shown to correlate with bacterial lung clearance (30). It has been reported that alveolar macrophages are essential for the rapid clearance of many pulmonary pathogens including *PA* (31), and a future direction may be to examine the effects of BOS-318 on clearance of chronic *PA* lung infections, which has many key clinical implications Great effort is taken to eradicate *PA* in pediatric CF patients after initial colonization, which requires treatment for months with two or more IV antibiotics. Post-lung transplant patients colonized with *PA* in the sinuses undergo lifelong nebulized antibiotics to keep these pathogens at bay and out of the transplanted organs. Finally, while CFTR modulators improve mucociliary clearance and reduce inflammation, these therapies do not eradicate *PA* (32).

On the contrary, depletion of alveolar macrophages has been shown to increase pulmonary neutrophil infiltration, tissue damage, and sepsis (33). Interestingly, we observed a downregulation of *Cyp1a1* RNA in the lungs of BOS-318-treated mice. When overexpressed in macrophages, this enzyme can impair phagocytosis of bacteria and increase mortality in an *E. coli* model of intraperitoneal sepsis (34). Pharmacologic *Cyp1a1* inhibition improved survival and bacteria clearance of mice in sepsis (34). Cyp1a1 activation may also suppress the cytotoxic effects of NK cells, particularly in anti-tumor immunity (35). Three main hurdles in developing *Cyp1a1* inhibitors has been selectivity and off-target effects, drug-drug interactions due to the common CYP pathway for metabolism, and membrane permeability which are key to reach intracellular targets. BOS-318 is highly selective, membrane-permeable, and may allow for impacting the downstream *Cyp1a1* pathway and bypassing these obstacles. Thus, furin inhibition could augment bacterial clearance by increasing AM phagocytosis and modulating *Cyp1a1* expression, although more studies are needed to understand the mechanisms in which furin inhibition refines alveolar macrophages phagocytosis and NK cell cytotoxicity.

In our model, furin inhibition strongly decreased chemoattractant MCP-1 concentration in BAL, which corresponds with attenuated neutrophil and NK cell recruitment. It has been observed that MCP-1 promotes NETosis in neutrophils *in vitro* (36). NK cell recruitment has been reported to be dependent on MCP-1 and critical for clearance of lung pathogens, such as aspergillus (37). As reported by Eleman *et al*., NK cells could be helpful or harmful in PA infection (38), and more studies are needed to characterize these immune pathways. In a pneumo-toxin mouse model, Exo-A-induced NK cell recruitment is reported to induce liver damage (39). In COPD, NK cells may drive lung tissue destruction through cytotoxicity against lung epithelial cells (40, 41). In a pulmonary ischemia-reperfusion model (IRI), Calabrese *et al*. observed an increase of NK trafficking from peripheral reservoirs into the lung tissue, and antibody-mediated NK cell depletion improved IRI-induced acute lung injury (42). NK cell depletion has also been shown to increase neutrophil recruitment in the lungs and mortality without impacting bacterial burden, suggesting a role for NK cells in modulating inflammation beyond bactericidality (43). Our work suggests that furin inhibition could reduce neutrophil and NK cell infiltration as well as their associated damages by enhancing alveolar macrophage-mediated bacterial clearance. More work is needed to fully determine the importance of furin in NK cell biology.

While our results are promising, several limitations need to be acknowledged. First, our study relied on murine models of Exo-A and PA-mediated acute lung injury, which may not fully recapitulate the acute-on-chronic, complex nature of infections observed in patients with CF or COPD. However, Douglas *et al*. reported that BOS-318 enhanced hydration and mucociliary transport (MCT) in human CF epithelial cells (14, 44). Our mouse models were in wild-type mice, and future studies could utilize CF mouse models despite known limitations in capturing structural human CF lung disease. Second, our approach with furin inhibition was done at the same time as infection, and future studies could attempt a rescue approach after bacterial pneumonia is established. Moreover, translating these results into chronic infection models will be critical to understand the long-term benefits and safety of furin inhibition such as CF (45, 46). Our hope is that BOS-318 could be a therapeutic option in patients with chronic *PA* lung infection as well as acute *PA* lung infection, which commonly trigger CF and COPD exacerbations. Finally, while BOS-318 is highly specific for furin, furin cleaves numerous substrates, which raises the question of specificity. Even if furin inhibition is safe and promotes resolution of PA acute lung infection, some studies suggest that furin inhibition could disrupt embryogenesis and homeostasis (47, 48). Thus, future studies employing lineage-specific furin deficiency models (e.g., alveolar macrophages, NK, or epithelial cells) may help clarify individual immune cell contributions during bacterial infection. Attention should be paid also in other cell types that could be important during acute and/or chronic infection such as endothelial cells. Indeed, silencing furin in mice resulted in reduced processing of pro-inflammatory and pro-fibrotic cytokine precursors endothelin-1 and TGF-β1 (49). Moreover, furin inhibition using previously described Hexa-D-arginine attenuated TGF‐β1‐induced fibroblast overproliferation and transmigration, and reduced pulmonary fibrosis in mouse model, suggesting that BOS-318 could also be used to treat airway fibrosis and remodeling like in COPD, CF and also idiopathic pulmonary fibrosis (27).

In summary, our study provides compelling evidence that furin inhibition using BOS-318 not only protects against Exotoxin A-mediated lung injury but also modulates key inflammatory pathways to enhance bacterial clearance. These findings pave the way for the use of protease inhibitors as new therapeutic strategies in the treatment of PA lung infections.

## AUTHOR CONTRIBUTIONS

OB designed and conducted experiments, analyzed the data, and wrote the manuscript. M.K performed experiments and analyzed the data, MRL provided resources and edited the manuscript, M.M performed experiments, analyzed the data, provided intellectual input and edited the manuscript. M.A.Y provided intellectual input, provided resources and materials and edited the manuscript.

## ACKNOWLEDGMENT

We thank the UCSF Parnassus Flow CoLab, RRID:SCR_018206 and DRC Center Grant NIH P30 DK063720 for using their LSRII flow cytometer “Kermit”, and the UCSF Genomic CoLab for the bulk RNA_seq run and analysis. This work was supported by the UCSF Physician Scientist Scholars Program (5014-138401-2015664-45), the Cystic Fibrosis Foundation LeRoy Matthews Award (YU18L0) and NIAID grant RO1 AI160167.

---

**Fig 1 supp:**
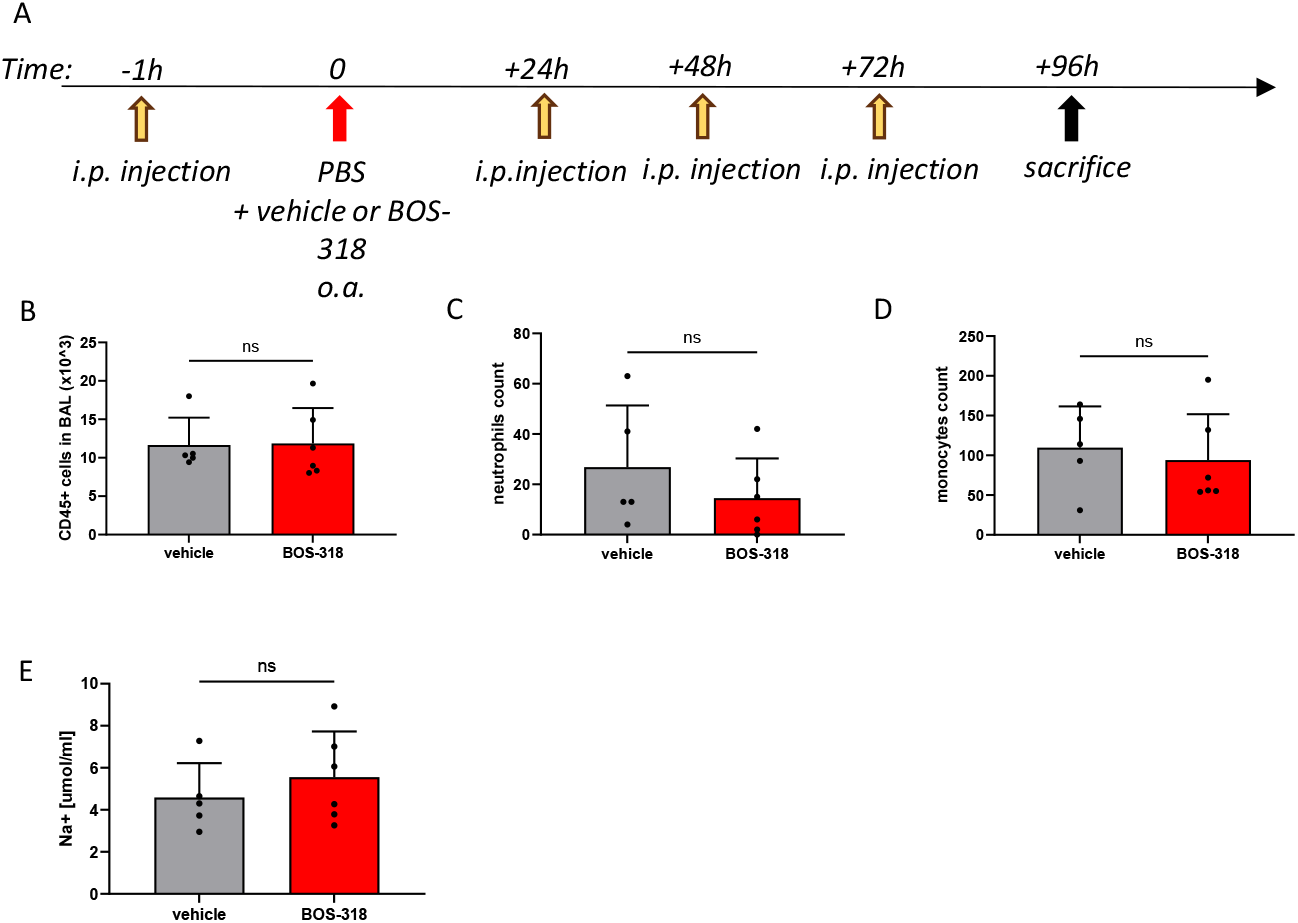
Furin inhibition doesn’t affect lung homeostasis. (A) Schematic depicting experimental procedure. Mice were injected with vehicle or furin inhibitor Ii.p. one hour prior oropharyngeal aspiration (o.a.) of PBS + vehicle or furin inhibitor, then animals were injected i.p. the 3 following days, then sac at 96h. (B-D) CD45+ cells, monocytes and neutrophils in BAL counted by flow cytometry. (n = 5-6 mice/group). (E) Sodium levels in BAL. (n = 5-6 mice/group) Unpaired t-test was used for statistics analysis.

